# How prolonged expression of Hunchback, a temporal transcription factor, re-wires locomotor circuits

**DOI:** 10.1101/556209

**Authors:** Julia L. Meng, Zarion D. Marshall, Meike Lobb-Rabe, Ellie S. Heckscher

## Abstract

In many CNS regions, neuronal birth timing is associated with circuit membership. In Drosophila larvae, we show U motor neurons are a temporal cohort—a set of non-identical, contiguously-born neurons from a single neuronal stem cell that contribute to the same circuit. We prolong expression of a temporal transcription factor, Hunchback, to increase the number of U motor neurons with early-born molecular identities. On the circuit level, this expands and re-wires the U motor neuron temporal cohort. On the cell biological level, we find novel roles for Hunchback in motor neuron target selection, neuromuscular synapse formation, dendrite morphogenesis, and behavior. These data provide insight into the relationship between stem cell and circuit, show that Hunchback is a potent regulator of circuit assembly, and suggest that temporal transcription factors are molecules that could be altered during evolution or biomedical intervention for the generation of novel circuits.

## Introduction

In many CNS regions, in many species, there is an association between neuron birth timing and neural circuit membership [1-13]. The mechanisms underlying this association are likely to be varied and are still poorly understood.

In complex brains, neurons are born from pools of neuronal stem cells. As stem cells divide, several factors are simultaneously changing each of which could influence the circuit membership of the resulting neuron. One factor is a dynamic environment—earlier-born neurons and later-born neurons can encounter different physical substrates or receive different signaling cues. A second type of factor is that earlier-born neurons have more time for processes like migration and axon extension. Third, over time, within the nucleus of a neuronal stem cell, the chromatin landscape can change. In addition, earlier-born and later-born neurons have different molecular profiles. Determining the precise role of each of the factor associated with birth time is crucial for understanding how circuits assemble.

Temporal transcription factors are molecules that are dynamically expressed in neuronal stem cells [14], and they have been suggested to play a role in circuit wiring [7]. The defining feature of a temporal transcription factor, as opposed to a cell fate determinant is that temporal transcription factors act in multiple lineages to specify the identity of neurons born at a certain time (e.g., early-born) regardless of cell type (e.g., motor neuron, interneuron, glia) [15]. Although temporal transcription factors play an important role in establishing the molecular diversity of neurons within a lineage, their role in regulating neuronal morphology, synaptic connectivity, circuit wiring, and behavior are far less understood [16]. Temporal transcription factors have also been called temporal identity factors, and the use of the term *<u>identity</u>* is often taken to mean that these factors are upstream of all of a neuron’s post-mitotic features [17,18], which is at the heart of the assumption that temporal transcription factors can regulate circuit wiring. However, data to support this assumption are limited [15].

Here, we probe the role of temporal transcription factors in circuit wiring by manipulating the expression of Hunchback (Hb) in the Drosophila motor system. Hb is one of the first-identified, and best-characterized temporal transcription factors [19]. The vertebrate homolog of Hb, Ikaros acts as a temporal transcription factor in the cortex and retina [20,21]. Although Hb is active in most Drosophila nerve cord neuroblasts, we focus specifically on the role of Hb in NB7-1 because Hb has been studied intensely in the NB7-1 lineage [19-23].

First, we characterize the axonal and dendritic morphologies of the first-born progeny of NB7-1, the U motor neurons. We show each U motor neuron makes a distinctive neuromuscular synapse on to a unique muscle located in the synergistic group of dorsal muscles. This shows that U motor neurons are an example of a recently-discovered, developmentally-based unit of circuit organization, a “temporal cohort” [24]. A temporal cohort is a block of non-identical neurons that have shared circuit membership (e.g., process similar sensory modality, project to synergistic muscles) that are contiguously-born from a single neuronal stem cell. The identification of the U motor neuron temporal cohort adds support to the idea that temporal cohorts are a common organizational unit in the Drosophila motor system.

Hb is normally expressed before the first two divisions of NB7-1 [19]. Here, we prolong Hb expression in NB7-1 for the entire duration of embryonic neurogenesis. On the circuit level, this expands and alters the wiring of the U motor neuron temporal cohort, resulting in a temporal cohort that should carry out a novel circuit level computation. On the cell biological level, we show Hb can completely control some mature motor neuronal features—nerve cord exit and axonal trajectory, but can incompletely control other features—neuromuscular synapse formation and dendrite morphogenesis. This shows that although Hb can control many mature neuronal features, other factors are also required. Our findings provide insight into the relationship between stem cell and motor circuit, parse the contributions of birth time related factors in circuit assembly, and suggest that temporal transcription factors are molecules that could be altered during evolution or biomedical intervention for the generation of novel circuits.

## Results

### The NB7-1 lineage contains a temporal cohort of U motor neurons

In this study, we probe the role of temporal identity factors in circuit wiring by manipulating the expression of Hunchback (Hb) in a single neuronal stem cell lineage, the NB7-1 lineage. Specifically, we focus on a set of five U motor neurons (U1-U5) produced by the first five divisions of NB7-1.

To lay a foundation, we characterized the circuit membership of the U motor neurons. In the Drosophila motor system, motor neurons innervate three anatomically and functionally distinct groups of muscles—dorsal, ventral, and transverse—each of which is driven by a separate neuronal circuit (Figure 1A-B) [25,26]. U motor neurons were thought to project to dorsal muscles, as well as other muscles, but their exact muscle targets were unknown [27-29]. To map muscle targets, and therefore circuit membership, we labeled single U motor neurons, and visualized their axons in embryonic stages as well as their neuromuscular synapses at larval stages. At both stages, U motor neurons can be identified by their characteristic position in the CNS and expression of the homeobox transcription factor, Even-skipped (Eve, Evx1/2 in vertebrates, Figure 1C-H). Although Eve is expressed by neurons in other lineages, it is exclusively expressed by U motor neurons in the NB7-1 lineage. In embryos, we find every U motor neuron axon projects to a unique, dorsal muscle (Figure 1D’-H’). In late-stage (L3) larvae, U motor neurons form neuromuscular synapses on dorsal muscles (Figure 1I, S1A). Specifically, the embryonic U1 motor neuron projects to Muscle 9 and is the larval motor neuron MN9-1b. Note, larval motor neurons are individually named based on muscle target and synaptic morphology, and so for example, MN9-1b is a <u>m</u>otor <u>n</u>euron that targets Muscle <u>9</u>, and has type <u>1b</u> (big) synaptic boutons. U2 projects to Muscle 10 and is the larval MN10-1b. U3 is MN2-1b, U4 is MN3-1b, and U5 is MN4-1b. All U motor neurons are excitatory, and we therefore conclude that all U motor neurons are members of the circuit responsible for driving contraction of dorsal muscles.

**Figure 1.**
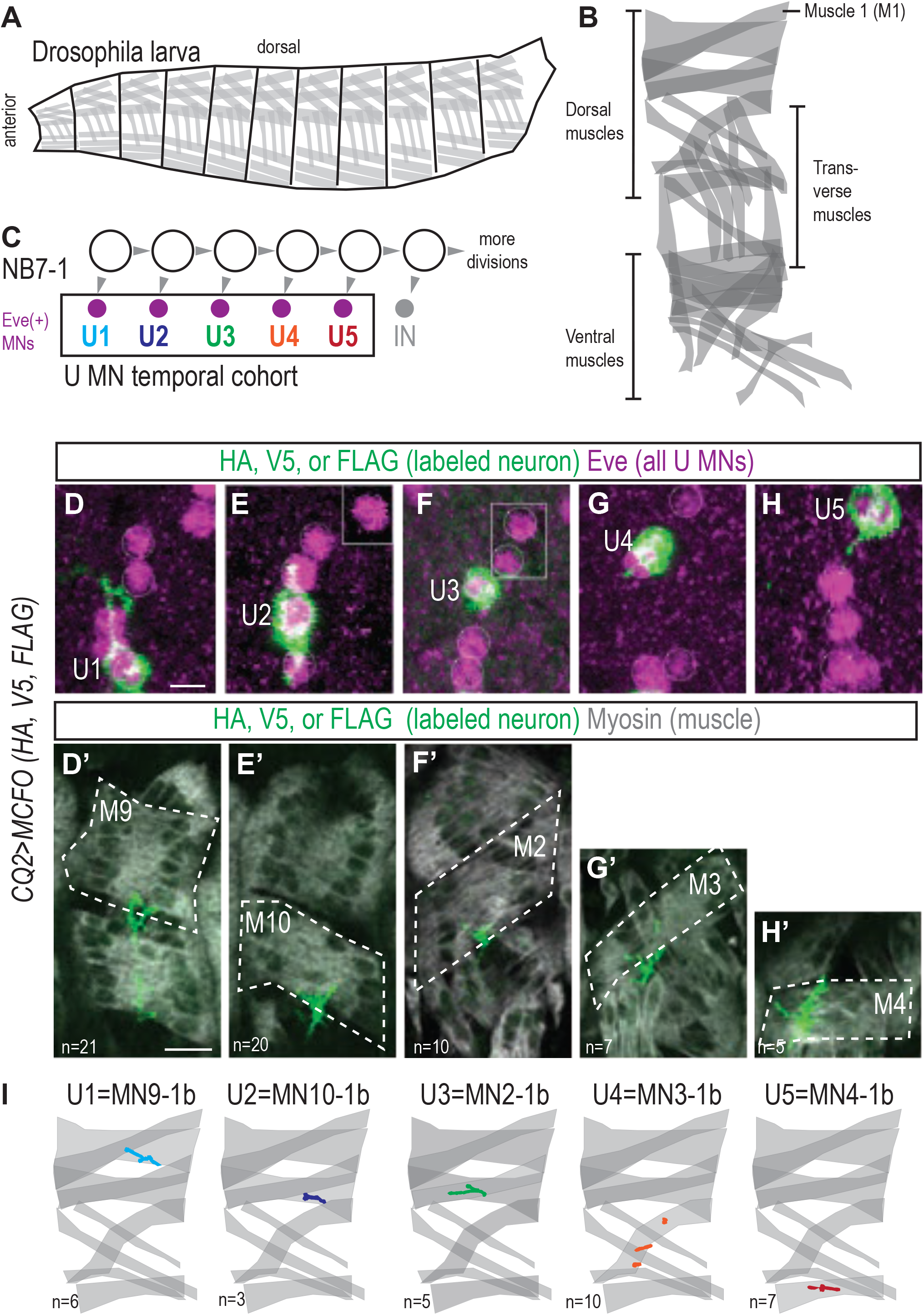
The NB7-1 lineage contains a temporal cohort of U motor neurons. (A-B) Illustrations of Drosophila larval muscles. The larval body is organized into repeated left-right mirror image hemisegments (A). In a hemisegment, individual muscles can be identified by characteristic morphology and have unique names (e.g. “Muscle 1”). Dorsal and ventral (oriented from anterior to posterior), and transverse (oriented from dorsal to ventral) muscle groups share innervation, inputs, and function, which is unique to each group. We therefore consider muscles within a group to be synergistic and controlled by a unique circuit (B). (C) Illustration of the divisions of NB7-1. Each gray arrowhead represents a cell division. MN is motor neuron and IN is interneuron. U MNs express the homeobox transcription factor, Even-skipped (Eve, Evx1/2 in vertebrates). A temporal cohort is a set of non-identical, contingously-born neurons from a single neuronal stem cell that contribute to one circuit. (D-H) Images of individually labeled U motor neuron cell bodies in the CNS of late stage embryos shown in ventral view. Multi-Color Flip Out (MCFO) transgenes were used to stochastically label neurons within a GAL4 pattern with membrane tethered epitope tags (HA, V5, FLAG). In embryos, MCFO transgenes were driven with a U MN-GAL4 line, *CQ2-GAL4 (hsFLP; CQ2-GAL4/+; UAS(FRT.stop)myr∷smGdP-HA, UAS(FRT.stop)myr∷smGdP-V5-THS-UAS(FRT.stop)myr∷smGdP-FLAG/+).* A ventral view is shown. Boxes are insets from a different focal plane. (D’-H’) Images of individually labeled U motor neuron from D-H are shown in lateral view. Axons project to unique dorsal muscles (dashes, M stands for muscle, e.g. M9 is Muscle 9). n = number of single-labeled Eve(+) cells. Midline is down. (I) Illustration of individual U MN neuromuscular synapses onto dorsal muscles in larvae. Embryonic (e.g., U1) and larval (e.g. MN9-1b) motor neuron names are above neuromuscular synapse location. n = number of single-labeled Eve(+) cells. Color code as in C. For an example of image data see Figure S1. Images in (D-H’) are shown with dorsal up, anterior to the left, Scale bars represent 5 microns and 10 microns (D’).

These data demonstrate that U motor neurons fulfill the criteria to be considered a “temporal cohort”—a block of non-identical, contiguously-born neurons from a single neuronal stem cell (Figure 1C) that share circuit membership (e.g., synergistic muscles, Figure 1I).

### Prolonged expression of Hb in NB7-1 generates extra Eve(+) motor neurons that project to dorsal muscles in embryos

Characterization of U motor neuron-to-muscle connectivity provides a system in which to ask how manipulation of temporal transcription factors impact circuit wiring. Temporal transcription factors have been suggested to play a role in circuit wiring, but this has not been demonstrated [7]. To test this idea, we prolong in NB7-1 the expression of the temporal identity transcription factor, Hunchback (Hb). In wild type, Hb is expressed during the first two NB7-1 divisions, and without Hb, motor neurons expressing molecular markers characteristic of U1 and U2 are absent [19]. The expression of Hb in NB7-1 has been extended in many different ways. In general this has two effects—it generates extra Eve(+) cells, and it alters the expression of U motor neuron molecular markers [19-21,30]. Depending on the specific manipulation, extra Eve(+) cells may have U1- and U2-like molecular identities, or other U-like molecular identities. Extra Eve(+) cells produced when Hb expression is prolonged have been suggested to be motor neurons [20]. Thus, an attractive, but untested hypothesis is that prolonged expression of Hb in NB7-1 can expand and re-wire the U motor neuron temporal cohort.

We manipulated Hb expression using a GAL4 line that drives specifically in NB7-1, “*NB7-1-GAL4” (gooseberry-DBD, achaete-VP16,* see Figure S2A-B) [21]. We use lineage specific manipulation because it allows us to rule out possible NB7-1 lineage non-autonomous effects of Hb, which is especially important when thinking about circuit wiring. We drove Hb using two copies of a *UAS-Hb* transgene, which we refer to as “NB7-1>Hb”. In NB7-1>Hb embryos, Hb expression in NB7-1 is prolonged and extra Eve(+) cells are generated (Figure 2A-C, S2C-D).

**Figure 2.**
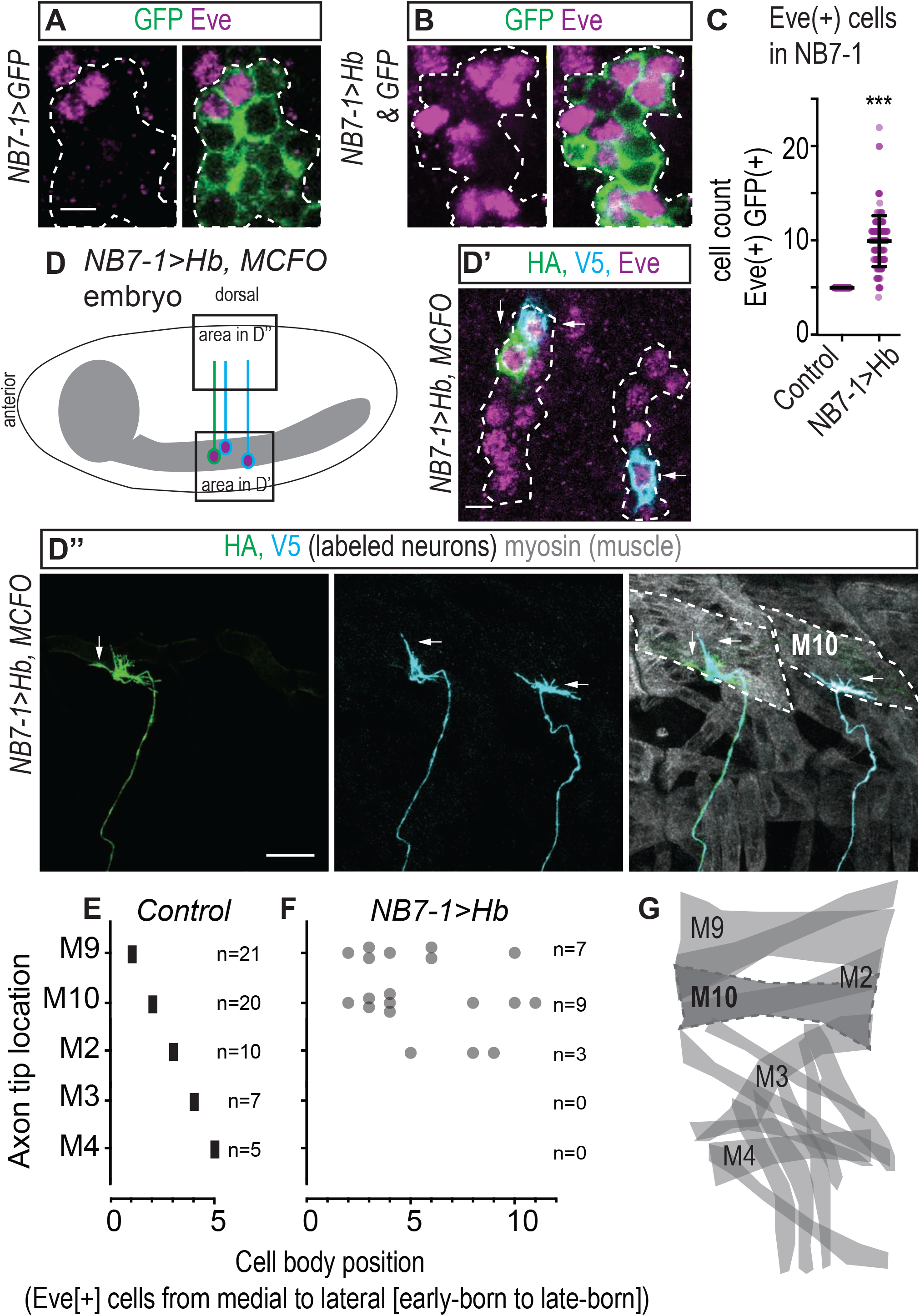
In NB7-1>Hb, extra Eve(+) cells are MNs that project to dorsal muscles. (A-B) Images of NB7-1 Eve(+) cells in the embryonic CNS. *NB7-1-GAL4* was used to drive GFP (*NB7-1-GAL4/+; UAS-myr-GFP/+*) or both GFP and Hb (*NB7-1-GAL4/UAS-Hb; UAS-Hb/UAS-myr-GFP*). NB7-1 cells are outlined (dashes). Prolonged expression of Hb in NB7-1 generates extra Eve(+) cells in embryos. (C) Quantification of Eve(+) cells in the NB7-1 lineage of Control and NB7-1>Hb embryos. Each dot represents number (Control n = 100, NB7-1>Hb n = 148) in an abdominal hemisegment. Average and standard deviation are overlaid. Unpaired t-test. ‘***’ p<0.0001. (D) Illustration of the embryo in D’ and D’’ shows that in NB7-1>Hb Eve(+) cells send axons to dorsal muscles. CNS is in gray. (D’-D’’) Images of a late stage, whole-mount, NB7-1>Hb embryo shown in ventral (D’) and lateral views (D’’). Individual Eve(+) cells are labeled with MCFO (*hsFLP; NB7-1-GAL4/ UAS-Hb; UAS(FRT.stop)myr∷smGdP-HA, UAS(FRT.stop)myr∷smGdP-V5-THS-UAS(FRT.stop)myr∷smGdP-FLAG/UAS-Hb*). (D) There are >5 Eve(+) cells from NB7-1 in each hemisegment (dashes), three of which are labeled. (D’’) Axons from Eve(+) cells project to Muscle 10 (M10, dashes). Vertical and horizontal arrows point to the same HA(+) and V5(+) cells, respectively (D’-D’’). (E-F) Quantification of individually-labeled Eve(+) cells. Position of the axon tip (muscle target) is plotted against cell body position (order from midline of Eve(+) cells). (E) Each box represents many cells. The number of cells is given as n=value at right. Data from Figure 1D-H’ are plotted. (F) Each circle represents a single cell. (G) Illustration of dorsal and transverse muscles with wild type U motor neuron target muscles labeled (M = Muscle). Dashes outline Muscle 10. Images are shown with dorsal up, anterior to the left. Scale bar is 5 microns (A, D’) and 10 microns (D’’).

In NB7-1>Hb larvae, we asked to what extent are the extra Eve(+) cells indeed motor neurons. In Drosophila, motor neurons have cell bodies in the CNS that send axons out of the nerve cord, which distinguishes them from interneurons whose axons do not leave the CNS, and from sensory neurons that have cell bodies in the periphery and send axons into the CNS. In isolated CNSs of both Control and NB7-1>Hb, we labeled individual Eve(+) cells, and in every case (Control [n=29] and NB7-1>Hb [n=77]) axons from Eve(+) cells exit the CNS. We conclude, in NB7-1>Hb larvae, all Eve(+) cells are motor neurons.

We asked, in NB7-1>Hb embryos, to which muscles do Eve(+) motor neuron axons project. We found every Eve(+) motor neuron (n=19) sends an axon to one of the dorsal-most muscles within the dorsal muscle group. This includes Muscles 9, 10, and 2, but not 3 and 4 (Figure 2D-G). We note that even Eve(+) cells that are located lateral-most in the cluster of NB7-1 Eve(+) cells can send axons to the dorsal-most muscles (Figure 2F). Because cell body location is associated with neuron birth time, this suggests that even the latest-born Eve(+) cell in NB7-1>Hb can extend axons to Muscles 9, 10, and 2.

Taken together, we conclude in NB7-1>Hb embryos, all Eve(+) motor neuron axons selectively project to the dorsal-most subset of dorsal muscles independent of neuronal birth time. These data support the idea that prolonged expression of Hb in NB7-1 could expand and re-wire the U motor neuron temporal cohort.

### Prolonged expression of Hb in NB7-1 increases and re-distributes neuromuscular synapses in larvae

Prolonged expression of Hb in NB7-1 re-routes motor neuron axons in embryos, which raises the possibility that this could expand and re-wire the U motor neuron temporal cohort. However, in both Drosophila and vertebrate motor systems, there is a complex relationship between axon pathfinding and synapse formation. In the Drosophila motor system, axon mistargeting often leads to delayed, but otherwise normal synapse formation [31-34]. In the vertebrate motor system, even though multiple axons reach a given muscle, only one actually forms a stable synaptic connection, whereas the others are eliminated via competition [35,36]. Here, we examine the extent to which prolonged expression of Hb in NB7-1 impacts motor neuron synapses on dorsal muscles at a late larval stage.

In wandering third instar (L3) larvae, we labeled all neuronal membrane including nerves and neuromuscular synaptic terminals (anti-HRP) (Figure 3A, E-F). Notably, this method allows us to look at the population of motor neurons, not just individually labeled neurons as described above.

**Figure 3.**
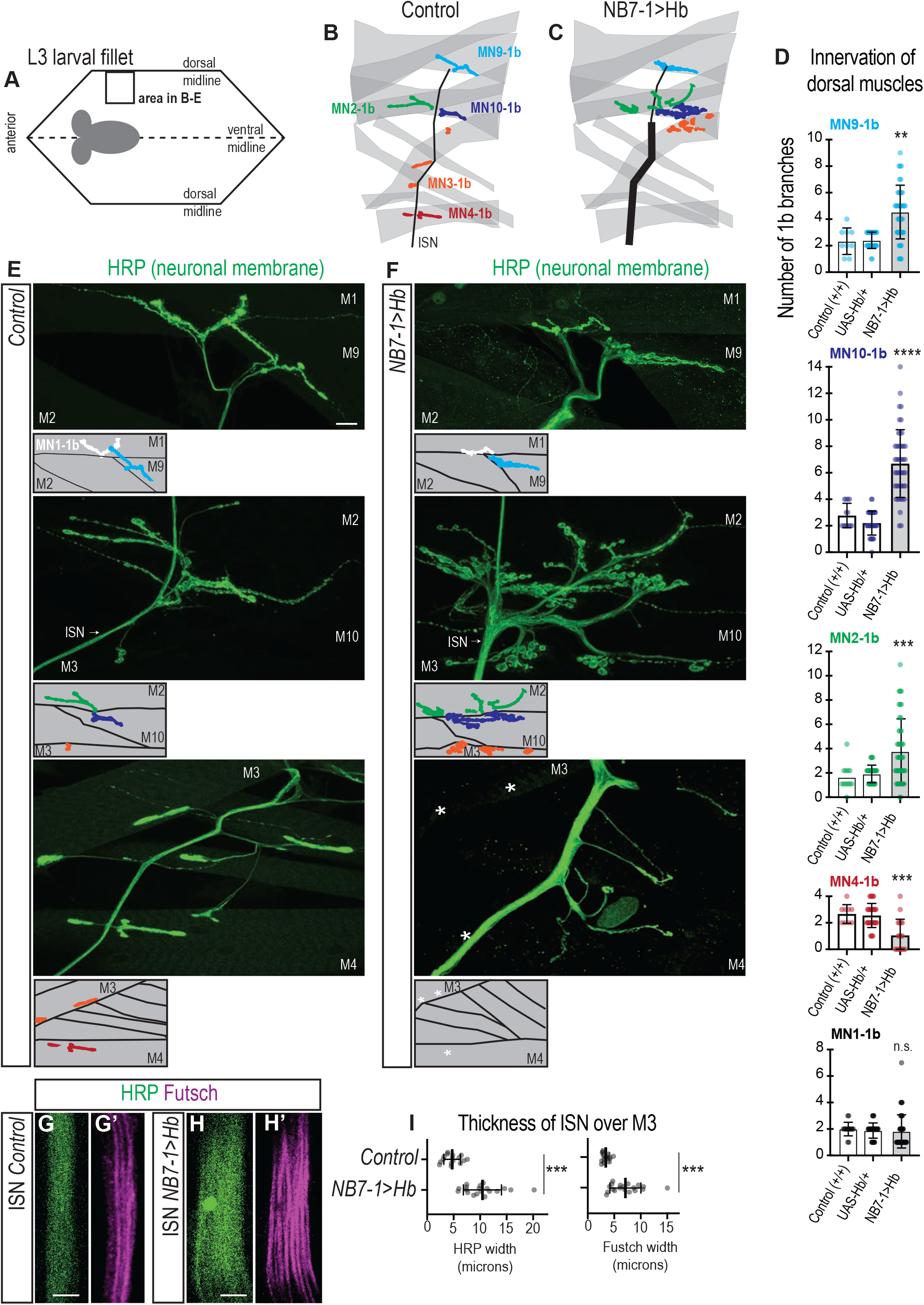
Prolonged expression of Hb in NB7-1 alters neuromuscular synapse on dorsal muscles in larvae. (A) Illustration of a Drosophila larval fillet preparation is shown with ventral midline dashed and CNS in gray. The position of dorsal muscles in one hemisegment is boxed. (B-C) Illustrations of neuromuscular synapses on dorsal muscles. In comparison to Control, in NB7-1>Hb, there is an increase in type 1b neuromuscular synapses branches on Muscles 9 (light blue), 10 (dark blue) and 2 (green), and loss of branches from Muscle 4 (red). The position of branches on Muscle 3 is shifted (orange). The intersegmental nerve (ISN, black line) contains more axons and is thicker. (D) Quantification of type 1b branch number on individual dorsal muscles. Control (+/+) (*w11118*), UAS-Hb/+ (*UAS-Hb/+; UAS-Hb/+)* NB7-1>Hb (*NB7-1-GAL4/UAS-Hb, UAS-Hb/+*). NB7-1>Hb is shaded. Note MN1-1b is a control muscle that lacks U MN innervation. (E-F) Images of neuronal membrane—both axons and neuromuscular synapses—on dorsal muscles in Control (E) and NB7-1>Hb (F). Arrow indicates ISN crossing M3 (see G-H for more). * indicates missing synapses. Below each image is a schematic of the image with the color scheme described in B-C. (G-H) Images of the ISN as it passes over M3 labeled both by a membrane marker (HRP) and a microtubule binding protein (Futsch). In NB7-1>Hb (H-H’), in comparison to Control (G-G’), the ISN is thicker. Note there is one Fustch-labeled microtubule bundle per axon (See Figure S1). (I) Quantification of ISN thickness. Image data is shown dorsal up anterior to the left. Scale bars represent 10 microns (E) and 5 microns (G-H). Quantification data is shown with each dot representing a value in an abdominal hemisegment. In D, Control n = 9, 13, 12, 9, 16, UAS-Hb n = 32, 35, 33, 36, 36, NB7-1>Hb n = 39, 46, 44, 18, 46, numbers listed from MN9-1b to MN1-1b or top to bottom. In I Control n = 16, NB7-1>Hb n = 19. Average and standard deviation are overlaid. ANOVA, corrected for multiple samples (D). Un-paried t-tests (I). ‘ns’ not significant, ‘**’ p<0.05, ‘***’ p<0.001, ‘****’ p<0.0001

The intersegmental nerve (ISN), passing over Muscle 3, contains axons traveling from the ventrally-located CNS to the dorsal-most muscles. In NB7-1>Hb, in comparison to Control the width of the ISN is significantly thicker (Figure 3B-C, E-F, G-I). Moreover, we visualized stable microtubule bundles that run through each axon (anti-Futsch, Figure S1B), and again see an increase in ISN thickness in NB7-1>Hb (Figure 3G-I). This demonstrates that in NB7-1>Hb, dorsally-projecting motor neurons survive into late larval stages.

To characterize neuromuscular synapses, we counted the number of type 1b branches on dorsal muscles in NB7-1>Hb and Control. In NB7-1>Hb larvae, there are significantly more 1b branches on Muscles 2, 9, and 10, with the largest increase on Muscle 10 (Figure 3B-F). The number of 1b branches is reduced on Muscle 4. And often, on Muscle 3, the 1b branch location is altered from a ventral location to a dorsal location (Figure 3B-F). There is no change in 1b branch number on dorsal muscles that are not targets of U motor neurons (Figure 3B-F). We note that the location of extra neuromuscular synapses in larvae resembles the location of Eve(+) growth cones seen in embryos. We conclude that when Hb is supplied in the neuroblast, axons are not just transiently targeted to abnormal locations, but that targeting is stable over the duration of larval life.

In summary, in NB7-1>Hb, there is large increase in the number of 1b synaptic branches on dorsal muscles, and these neuromuscular synapses are not distributed across dorsal muscles in the same pattern as in Control. These data are consistent with the idea that prolonged expression of Hb in NB7-1 expands and re-wires the U motor neuron temporal cohort.

### In NB7-1>Hb larvae, extra synaptic arbors contain functional synapses

Because temporal cohort membership implies circuit function, not just circuit anatomy, we needed to understand the extent to which the extra synaptic branches in NB7-1>Hb contain functional synapses. To address this issue, we use fixed tissue staining, functional calcium imaging, and larval behavior.

In fixed tissue L3 fillets, we stained for a panel of synaptic marker proteins. There is normal abundance and localization of pre-synaptic markers—for active zones (Brp), stable microtubule loops (Fustch), and synaptic vesicles (Syn)—as well as post-synaptic markers—for post-synaptic density (DLG) and neurotransmitter receptors (GluRIIA) (Figure 4A-G). Thus, on the cell biological level, neuromuscular synapses in NB7-1>Hb are not different from Control.

**Figure 4.**
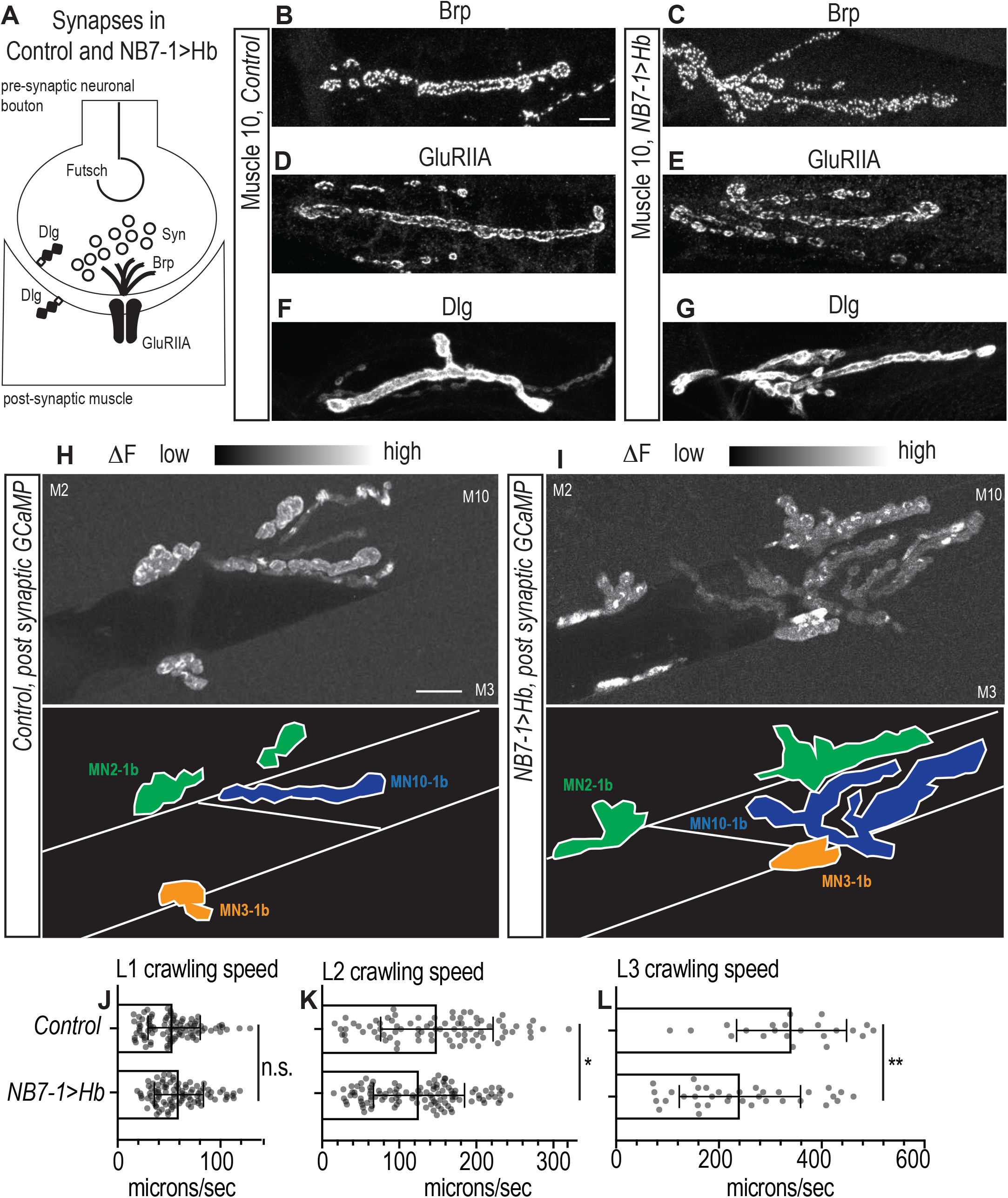
Altered synaptic arbors on dorsal muscles contain functional synapses. (A) Illustration of sub-cellular localization of neuromuscular synapse markers in Control and NB7-1>Hb. Futsch labels microtubules; Syn (Synapsin) labels neurotransmitter filled synaptic vesicles; Brp (Brunchpilot) labels vesicle release sites, or active zones; GluRIIA (Glutamate Receptor IIA) labels neurotransmitter receptors; Dlg (Discs large) a scaffolding protein is enriched in the post-synaptic density. (B-G) Images of neuromuscular synapses on Muscle 10. There is no difference in distribution or abundance of synaptic markers between Control (*w1118)* and NB7-1>Hb (*NB7-1-GAL4/UAS-Hb, UAS-Hb/+*). (H-I) Images of changes in fluorescence intensity of an calcium indicator for synaptic activity. GCaMP was targeted to the post-synaptic density (DLG in F-G). When pre-synaptic vesicles are released from active zones (Brp in B-C) post-synaptic neurotransmitter receptors respond (GluRIIA in D-E), increasing GCaMP fluorescence intensity (see Figure S3 for details). Top images show post-synaptic responses (delta F) in Muscles 2, 3, and 10 (M = muscle) in Control (*NB7-1-GAL4/+; MHC-CD8-GCaMP6f-Sh/+)* and NB7-1>Hb (*NB7-1 GAL4/UAS Hb; MHC-CD8-GCaMP6f-Sh/UAS Hb)* over a 5 minute imaging period. Bottom illustrations are tracings of 1b synapses. (J-L) Quantification of larval crawling speed. First, second, or third instar larvae (L1, L2, L3 respectively) were placed on agarose arenas and tracked for >1 minute. At each time point, larval centroid position was calculated, an approximation of the larval center of mass, and used to calculate larval crawling (centroid movement/time). Each dot represents the average speed for a Control (*w1118)* or NB7-1>Hb (*NB7-1-GAL4/UAS-Hb, UAS-Hb/+*) larva. Control n = 94, 82, 22, NB7-1>Hb n = 95, 114, 35 for L1, L2 and L3 respectively. Average and SEM are overlaid. Un-paired t-Test. ‘n.s.’ not significant, ‘*’ p<0.05, ‘**’ p<0.001. For images scale bars represent 10 microns (B, H).

We visualized neuromuscular synapse activity with a post-synaptically localized calcium sensor [37]. In L3 larval fillets, we imaged spontaneous release of individual synaptic vesicles from 1b branches on Muscles 1, 2, 9, and 10. In both Control and NB7-1>Hb, within a five-minute imaging period, we found at least one post-synaptic response in every 1b synaptic branch on every muscle imaged (Control [n = 6] and NB7-1>Hb [n =5] Figure 4H-I, S3). We conclude in NB7-1>Hb, extra arbors on dorsal muscles contain functional synapses.

To assess the degree to which locomotor circuits are functionally re-wired, we assayed larval crawling behavior. We placed larvae on agarose arenas and tracked over time the position of the larval centroid, an approximation of the center of mass. We calculated average crawling speed in Control and NB7-1>Hb in larvae at stages L1, L2, and L3. In L3 larvae, there is a significant decrease in crawling speed in NB7-1>Hb in comparison to Control (Figure 4L). Notably, this defect is detected at L2, but not at L1 (Figure 4J-K). This shows the phenotype emerges over time perhaps because L2 and L3 have increased larval crawling speeds in comparison to L1 (Figure 4J-K). These data are consistent with the idea that prolonging the expression of Hb in NB7-1 functionally re-wires locomotor circuits.

Together our anatomical (Figures 2-3) and functional (Figure 4) data provide strong support for the idea that prolonged expression of Hb in NB7-1 expands and re-wires the U motor neuron temporal cohort.

### In NB7-1>Hb, extra U motor neurons are born at abnormally late times

Having shown that Hb is able to expand the U motor neuron temporal cohort, we next asked, where do the extra U motor neurons come from? One model is that extra U motor neurons are generated by increased proliferation of the neuroblast. Indeed, Hb expression has been shown to regulate the cell cycle [38]. Alternatively, extra U motor neurons could be generated instead of other cell types in the lineage—either later-born neurons, or U siblings (neurons that are born from the same ganglion mother cell as U motor neurons, Figure 5A). Notably, the possibility that U motor neurons are generated at abnormally late times is consistent with the observation that providing pulses of Hb late in NB7-1 lineage development can generate extra U motor neurons [20,30]. Here, we directly test each possibility.

**Figure 5.**
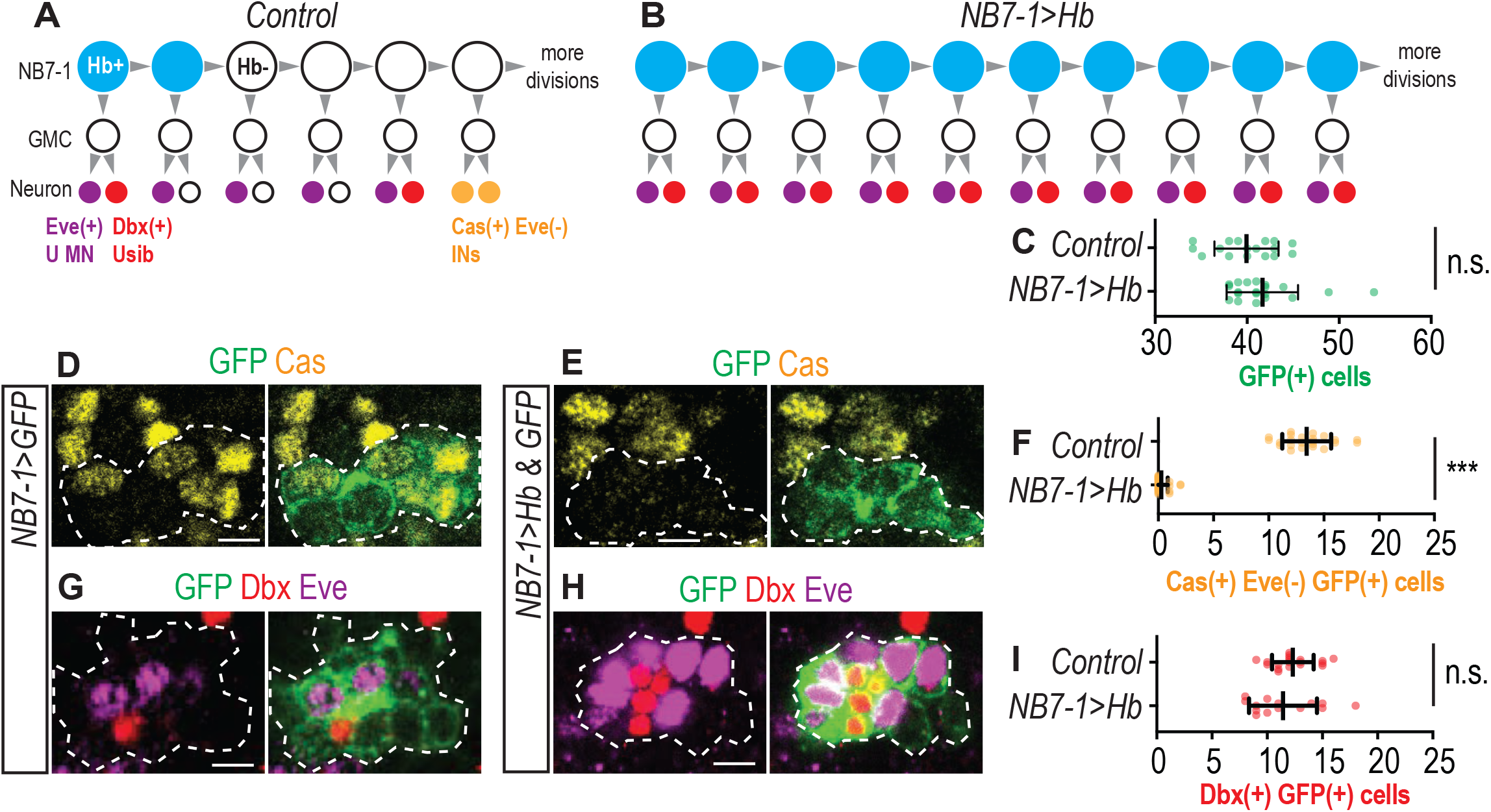
In NB7-1>Hb embryos, U motor neurons are produced abnormally late times in development. (A-B) Illustrations of lineage progression in Control and NB7-1>Hb. GMC stands for Ganglion Mother Cell, a transient cell that divides to generate different siblings (e.g., U motor neuron [MN] and U sibling [Usib]). In NB7-1>Hb, Hb is expressed during all NB7-1 divisions. Extra Eve(+) cells are produced when later-born NB7-1 interneurons (IN) are normally produced. (C) Quantification of neurons in the NB7-1 lineage in Control (*NB7-1-GAL4/+; UAS-myr-GFP/+*) and NB7-1>Hb (*NB7-1-GAL4/UAS-Hb; UAS-myr-GFP/UAS-Hb).* There is no significant difference in the total number of neurons in the NB7-1 lineage, and so increased proliferation cannot account for the increased number of U MNs. (D-E) Images, single confocal z-slices, of the embryonic CNS dorsolateral to U MNs. In wild type, Cas labels a set of later-born NB7-1 neurons (D), and in NB7-1>Hb virtually all Cas(+) cells are lost (E). (F) Quantification of Cas(+) Eve(-) INs from the NB7-1 lineage. (G-H) Images, single confocal z-slices, of the embryonic CNS at the level of U1 and U2. In wild-type, Dbx labels a small neuron from NB7-1 that is directly adjacent to U1, which the U1sib (G). In NB7-1>Hb, there is an increase both U1sibs and U MNs (H). (I) Quantification of Dbx(+) INs from the NB7-1 lineage. In addition to two Dbx(+) Usibs (REF) there more Dbx(+) cells in the NB7-1 lineage. The total number of Dbx(+) cells is not changed. For all image data anterior is up with the midline to the left. NB7-1 cells are dashed. Scale bars represent 5 microns. For quantifications each dot represents values in an abdominal hemisegment. Control n = 17, 23, 19, NB7-1>Hb n = 20, 30, 14, for C, F, I, respectively. Average and standard deviation are overlaid. Un-paired t-Test. ‘n.s.’ not significant, ‘***’ p<0.0001.

We tracked all NB7-1 progeny in Control and NB7-1>Hb embryos using a GFP reporter. First, we counted the total number of GFP(+) NB7-1 progeny, and find a slight increase in *NB7-1>Hb* in comparison to Control (Figure 5C), which is not enough to account for the number of extra U motor neurons. Second, we found that in wild-type, later-born NB7-1 interneurons express the Zinc-Finger transcription factor, Castor (Cas), but not Eve, and are located dorsolaterally to U motor neurons (Figure 5D) [19]. In NB7-1>Hb embryos, there is near complete loss of cells expressing Cas (Figure 5E-F). Third, in wild type, we found the Homeodomain transcription factor Dbx labels a subset of U sibling neurons, which are small, cells adjacent to U motor neurons, as well as other later-born interneurons in the NB7-1 lineage (Figure 5G) [39]. Using Dbx in NB7-1>Hb, we find no decrease in U siblings (Figure 5H), and so conversion of U siblings cannot account for the extra U motor neurons. However, overall there is no significant change in the total number of Dbx(+) neurons in the NB7-1 lineage (Figure 5I), suggesting the number of later-born Dbx(+) interneurons is reduced. Taken together, we conclude that in NB7-1>Hb, extra U motor neurons are generated at the time when neurons with later-born molecular identities (Cas[+]) should normally be generated (Figure 5B). Thus, prolonged expression of Hb in NB7-1 expands the size of the U motor neuron temporal cohort by causing later-born neurons to take on U motor neuron characteristics.

### In NB7-1>Hb embryos, a majority of U motor neurons have U1-like molecular identities

In NB7-1>Hb, the U motor neuron temporal cohort is re-wired because the distribution of neuromuscular synapses on dorsal muscles is different from Control. There are increased neuromuscular synapses on Muscles 2, 9, and 10, which in wild type are U1-U3 motor neuron muscle targets. The location of neuromuscular synapses on Muscle 3 (U4 target) are redistributed from the ventral to dorsal side of the muscle. There are fewer synaptic branches on Muscle 4 (U5 target). A simple explanation for the observed distribution of neuromuscular synapses in NB7-1>Hb larvae is that prolonged expression of Hb in NB7-1 generates early-born U motor neurons, but not late-born U motor neurons. At the heart of this explanation is the assumption that a neuron’s synaptic partner can be predicted by its embryonic molecular identity. And so, here, we characterize the embryonic molecular identities of U motor neurons in NB7-1>Hb.

Hb expression has been manipulated using a variety of different drivers (e.g., *hs-GAL4, En-GAL4, Pros-GAL4*, etc.), and each manipulation produces a set of Eve(+) cells with different embryonic U molecular identities [19-21,30]. We used combinations of molecular markers—Krupple, Zfh2, and Hb, Runt—to characterize U motor neuron identities in NB7-1>Hb (Figure 6A). Note, we use Hb in two capacities in this experiment. We manipulate Hb in the neuroblast, and we use Hb as marker for U1- and U2-like molecular identity because in wild type U1 and U2 motor neurons actively transcribe Hb [21]. To distinguish actively-transcribed Hb (an identity marker) from Hb inherited from neuroblast cytoplasm, we stained at late embryonic stages to allow for Hb protein turnover as in Pearson and Kohwi [20,21]. In NB7-1>Hb embryos, Eve(+) cells have U1-, U2-, and U3-like molecular identities, of these nearly 70% are U1-like. There are almost no Eve(+) cells with U4- or U5-like molecular identities (Figure 6B-E). We conclude that prolonged expression of Hb in NB7-1 using *NB7-1-GAL4* generates U motor neurons with early-born, but not late-born molecular identities.

**Figure 6.**
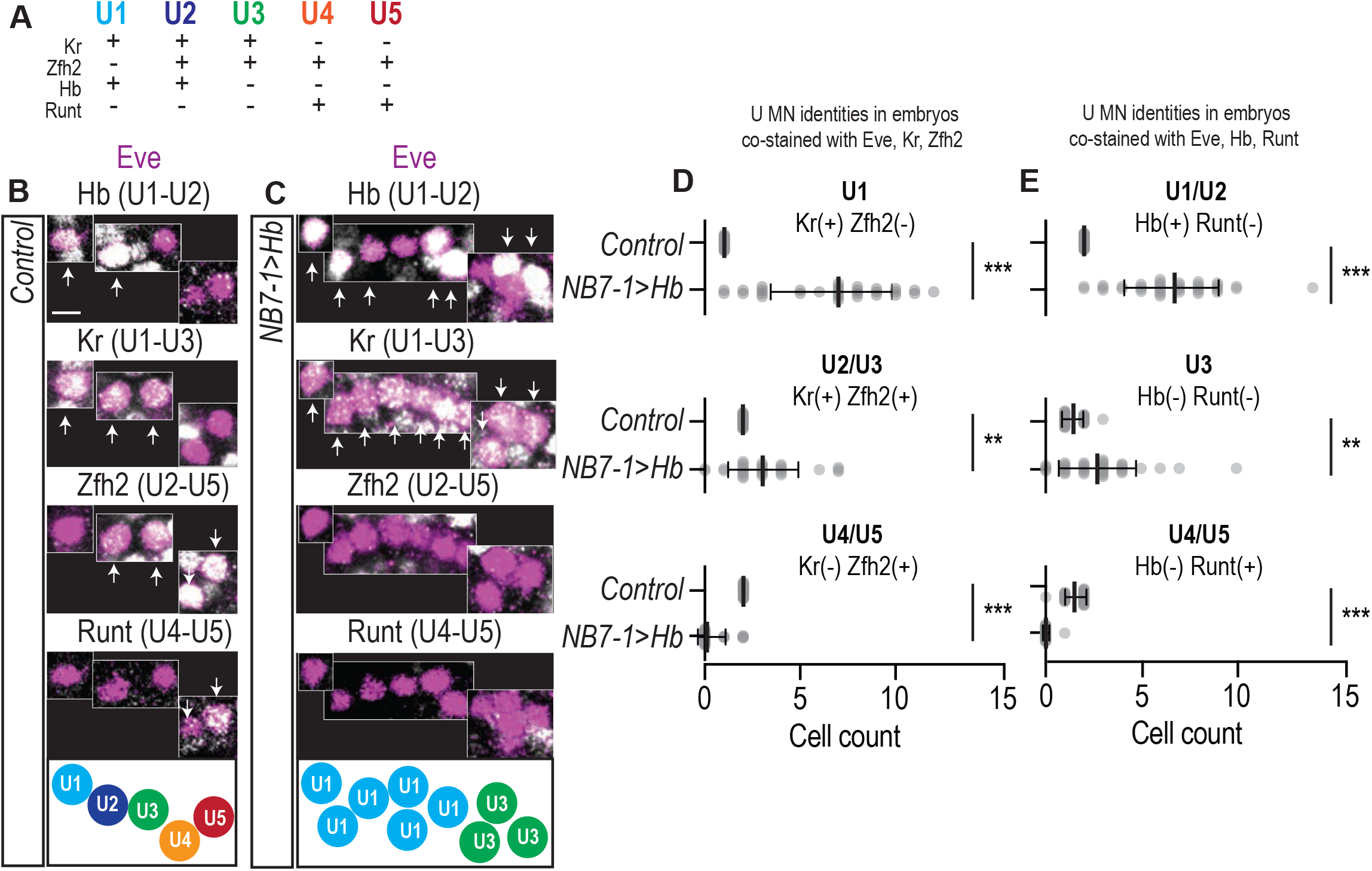
In NB7-1>Hb embryos, most U MNs have U1-like molecular identities. (A) Illustration of U MNs, which express distinct combinations of transcription factors—Kruppel (Kr), Zfh2, Hb, and Runt—that can be used to score embryonic U MN molecular identity. “+” indicates expression, and “-“ no expression. (B-C) Images of marker expression (white) in Eve(+) cells (magenta) in late-stage embryonic CNSs of Control (*w1118)* and NB7-1>Hb (*NB7-1-GAL4/UAS-Hb; UAS-Hb/+*). Extra extra Eve(+) cells are produced, which have U1-U3 molecular identities. Boxes from different z-planes. Arrows indicate co-expression. Anterior is up with the midline to the left. Scale bar is 5 microns. Summary illustrations of molecular identities shown at the bottom. (D-E) Quantification of marker gene expression. Each dot represents the cell count in a single abdominal hemisegment in late stage embryos. Control n = 25, 36, NB7-1>Hb n = 30, 38 in D and E, respectively. Average and standard deviation are overlaid. Un-paired t-Test. ‘n.s.’ not significant, ‘**’ p<0.0001, ‘***’ p<0.0001.

These data suggest that the observed distribution of neuromuscular synapses on dorsal muscles in NB7-1>Hb can be partially explained by embryonic U motor neuron molecular identity. However, these data argue against the assumption that a U motor neuron’s neuromuscular synaptic partner can be accurately predicted by its embryonic molecular identity. For example, in NB7-1>Hb embryos, the greatest increase in molecular identities is of U1-like neurons, yet in NB7-1>Hb larvae, the greatest increase in neuromuscular synapses is on Muscle 10 (U2 target). Furthermore, in NB7-1>HB embryos, there are no neurons with U4-like molecular identities, but Muscle 3 (U4 target) contacts are not missing, although they are abnormally located. We favor a model in which embryonic molecular identity can predict nerve root exit and axonal trajectory (Figure 2), but that the process of forming specific neuromuscular synaptic partnerships is not completely controlled by Hb.

### Prolonged expression of Hb in NB7-1 and other factors linked to neuronal birth time influence U motor neuron dendrite morphology

Given that prolonged expression of Hb in NB7-1 profoundly affects U motor neuron axonal trajectories, we also wondered if Hb influences dendrite morphology and therefore inputs onto U motor neurons. Of note, in the Drosophila motor system, dendrite morphogenesis and axonal targeting are not linked [26], and so, just because U motor neuron axons are altered by Hb expression does not imply there will be changes in dendrites.

One possibility is that Hb expression differentially affects axons and dendrites. In this scenario, in NB7-1>Hb, the U motor temporal cohort would contain “hybrid” motor neurons, which would map abnormal inputs onto dorsal muscles, change the computation carried by U motor neurons, and create a novel circuit level function for the U motor neuron temporal cohort. A second possibility is that Hb expression could affect both axons and dendrites similarly. In this scenario, extra U motor neurons would be replicas of wild type U motor neurons, and this would not fundamentally change the computation carried out by the neurons. To distinguish between these models, we characterized dendrite morphology in both Control and NB7-1>Hb larvae.

First, in Control, we characterized individual U motor neuron dendrite morphology in both embryos and larvae. At both stages, dendrites of U1 and U2 cross the midline, whereas dendrites of U3, U4, and U5 do not cross the midline (Figure 7A-B, S4). In Drosophila, neuronal cell body position is a proxy for neuronal birth time because there is little cell migration, and as a neuron buds off from its neuroblast, early-born progeny are pushed medially away from their laterally placed stem cell parent. In larvae, we measured the distance of each Eve(+) cell body from the midline as a proxy for birth time, and plotted cross/no cross dendrite morphology versus cell body position. In this plot, U motor neurons with cell bodies closest to the midline (early-born) have dendrites that cross the midline and U motor neurons with cell bodies more lateral (later-born) have dendrites that do not cross the midline (Figure 7D). These data show that in Control, correlated with neuronal birth time, U motor neurons have two morphologically distinct types of dendrites—those that cross the midline and those that do not.

**Figure 7.**
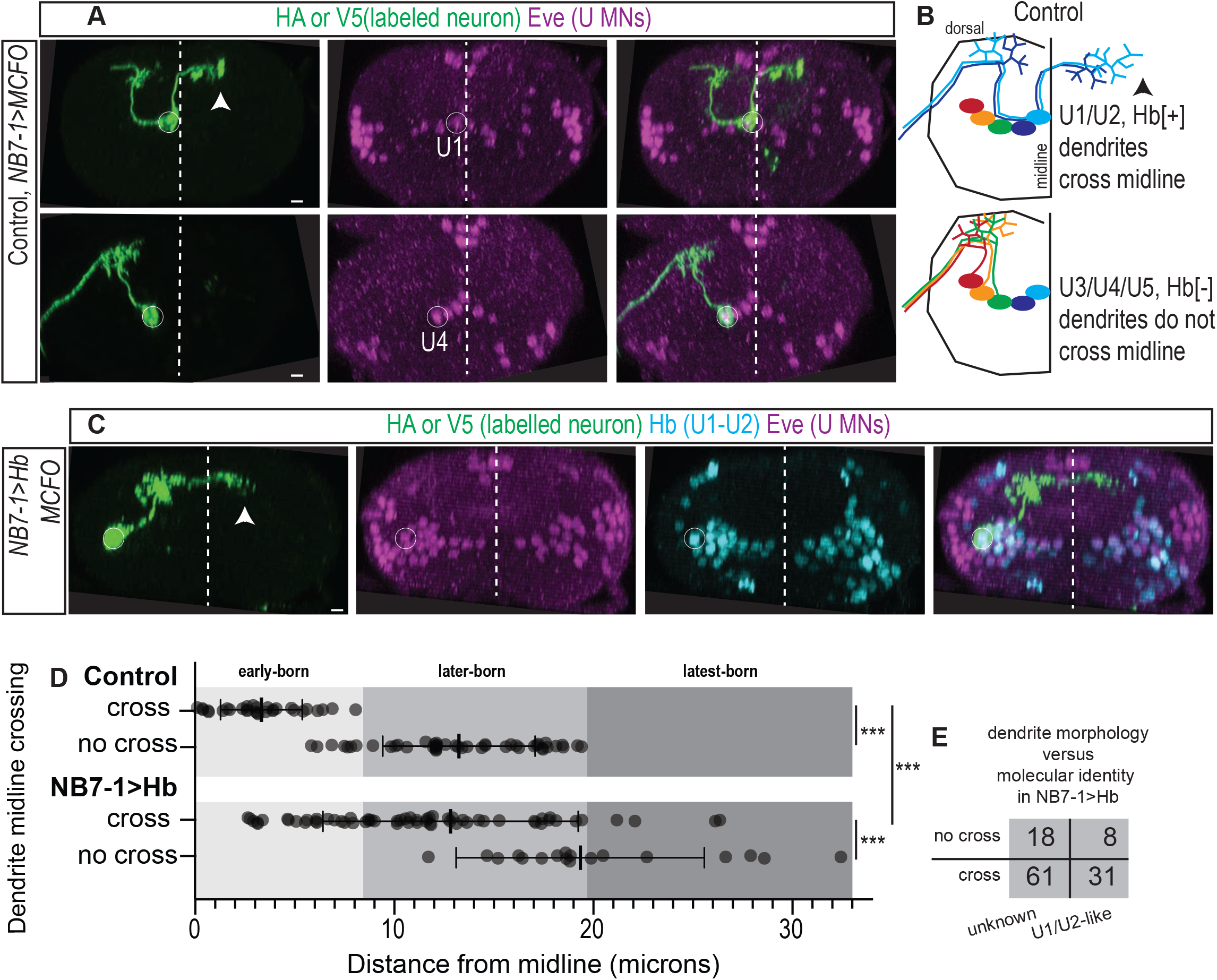
Prolonged expression of Hb in NB7-1 alters dendritic arborization. (A) Images of U MN dendrite morphology in the L1 CNSs of Control shown in transverse section. Individual U MNs were labeled (*hsFLP; NB7-1-GAL4/+; UAS(FRT.stop)myr∷smGdP-HA, UAS(FRT.stop)myr∷smGdP-V5-THS-UAS(FRT.stop)myr∷smGdP-FLAG/+).* U1 sends dendrites across the midline, whereas U4 does not. (See Figure S4). (B) Illustration of dendrite morphology, U motor neuron identity, and Hb expression in Control. Arrowhead points to U1/U2 neurons dendrites, which project across the midline. (light blue is U1, dark blue is U2, green is U3, orange is U4, and red is U5). (C) Images of U MN dendrite morphology in the L1 CNSs of NB7-1>Hb. Individual U MNs were labeled (*hsFLP; NB7-1-GAL4/UAS-Hb; UAS(FRT.stop)myr∷smGdP-HA, UAS(FRT.stop)myr∷smGdP-V5-THS-UAS(FRT.stop)myr∷smGdP-FLAG/UAS-Hb).* In this example, in NB7-1>Hb, a laterally-located, Hb(+) U MN sends a dendrite across the midline. (D) Quantification of individually labeled neurons in Control and NB7-1>Hb. Neurons were subdivided into groups based on dendrite midline crossing phenotype into “cross” and “no cross” groups, and plotted against cell body position (in microns from midline), which is a proxy for birth time. In Control, early-born neurons (light gray) have dendrites that cross the midline, whereas later-born (medium gray) neurons do not. This pattern is also seen in NB7-1>Hb, but shifted laterally (toward later-birth times). Average and standard deviation are overlaid. Ordinary one-way ANOVA, corrected for multiple samples. ‘***’ p<0.0001. (E) Quantification of individually labeled neurons in NB7-1>Hb. Neurons in the U1/U2-like category are labeled by Hb. Neurons in the unknown category were not co-stained for Hb and these data are also plotted in D. Images are shown with dorsal up, arrowheads pointing to dendrites crossing midline (dashed). The cell body location for labeled neurons are circled. Scale bar is 5 microns.

In NB7-1>Hb larvae, we conducted the same experiments and analysis, and found a majority of U motor neurons send dendrites across the midline (Figure 7C-D). The plot of dendrite morphology versus cell body position has a similar pattern to that seen in wild type, but shifted laterally (Figure 7D). This shows that in NB7-1>Hb, some U motor neurons that send dendrites across the midline are born at abnormally late times in comparison to Control. This is consistent with the idea that, at least for some neurons, Hb expression can affect both axons and dendrites.

In NB7-1>Hb, not all dendrites cross the midline. Moreover, dendrite morphology is associated with neuronal birth time (Figure 7D), which suggests that factors related to neuronal birth time influence dendrite morphology. These factors could include U1/U2-like molecular identities, or other factors (e.g., CNS neurogenesis, changing neuroblast chromatin landscape). To distinguish between these possibilities, in NB7-1>Hb we labeled, single neurons, and costained with the U1/U2 marker, Hb. In NB7-1>Hb, we found U1/U2-like Hb expression in many (n=39) U motor neurons. Of these Hb(+) neurons, a significant proportion (n=8) do not send dendrites across the midline (Figure 7E). These data show that in NB7-1>Hb larvae, birth time related factors influence dendrite morphology. Furthermore, because in NB7-1>Hb there is no association between birth time and axon target (Figure 2F) data in this section suggest that prolonged expression of Hb in NB7-1 generates some hybrid U motor neurons. Therefore, it is expected that prolonged expression of Hb in NB7-1 creates a novel dorsal muscle circuit configuration.

## Discussion

### The Drosophila motor system contains multiple different types of temporal cohorts

In the Drosophila nerve cord, we recently showed neurons from a single neuroblast can be organized into blocks of non-identical, yet functionally-related sets that contribute to different circuits. Specifically, early-born neurons in the NB3-3 lineage process mechanical sensory information and are members of an escape circuit, whereas late-born neurons in the NB3-3 lineage process proprioceptive cues and are members of a circuit that refines locomotion [24]. These blocks of lineage-related neurons are called “temporal cohorts”, defined as groups of contiguously-born neurons from a stem cell that have similar circuit level function.

Temporal cohorts are likely to be found in many lineages, brain regions, and organisms, and therefore could be fundamental, developmentally-based units of neuronal circuit organization. Despite the abundance of suggestive evidence for temporal cohorts, there are few explicit examples. For example, the vertebrate spinal cord may contain temporal cohorts—inhibitory Renshaw cells, which contribute to a motor neuron feedback circuit, and Ia interneurons, which contribute to a reflex circuit, are sequentially produced by neuronal stem cells within a small domains [40,41]. However, it is unknown the precise birth time or stem cell parents of these neurons. In other CNS regions (e.g., fly mushroom body and antennal lobe, vertebrate neocortex and retina), there are major transitions in lineage progression with stem cells producing dramatically different cell types over time [42-48], but the circuit membership of these neurons is poorly-defined at single neuron resolution. This illustrates why there are few explicit examples of temporal cohorts. To show that sets of neurons are *bona fide* temporal cohorts requires a large amount of information about individual neurons, including their stem cells parent, birth time, as well as a demonstration of circuit membership.

Here, we identify a temporal cohort in the NB7-1 lineage comprised of U motor neurons (Figure 1). These data demonstrate that temporal cohorts can be found in lineages other than NB3-3 and that temporal cohorts can be composed not just of interneurons, but also of motor neurons. Notably, the U MN temporal cohort innervates the dorsal muscles, which is just one of three muscle circuits in the Drosophila larval motor system (Figure 1A-B) [25]. This raises the question of whether other groups of muscles are innervated by other temporal cohorts. Nonetheless, the identification of the U motor neuron temporal cohort supports the idea that temporal cohorts are developmental units fundamental to circuit organization.

### Expanding our understanding of temporal transcription factors

Our data show that Hb is sufficient, at least partially, to control many post-mitotic neuronal features—nerve cord exit, axon trajectory, neuromuscular synapse formation, and dendrite morphology (Figure 2, 3, 4, 7). However, we note there are limits on Hb’s ability to control neuronal features and those limits are likely to include plasticity in neuromuscular synapse formation, and birth time related factors (e.g., dynamic CNS, intrinsic changes in the neuroblast).

Hunchback is a founding member of the class of transcription factors collectively termed, temporal identity factors, or temporal transcription factors. Although, first identified in the Drosophila motor system, temporal transcription factors have been found in many other Drosophila brain regions, and in vertebrates [16,49-54]. But the biology of temporal identity factors is incompletely understood. They have been shown to alter neuronal molecular markers, and some mature neuronal features including neurotransmitter identity and axonal trajectory. Recently, knock down of the temporal transcription factor, Eyeless was shown to affect neuronal morphology in adult fly central and affect navigational behavior [16]. Our study, re-enforces and broadens these findings. We also find impact on neuronal morphology and behavior, and do so using gain-of-function manipulation of a different temporal transcription factor in the motor system. Moreover, previous research has stopped short of characterizing synaptic partnerships and functional synapse formation. Our data therefor significantly extends our understanding of the biology of temporal transcription factors.

### Altered expression of a single transcription factor has the ability to re-wire circuits

The study of transcriptional control of circuit assembly is in its infancy. We have a firm understanding of how combinations of transcription factors work together to specify different neuronal types, assessed by marker gene expression, but a relatively poor understanding of how these specification events are related to mature circuit wiring.

Our data show that prolonged Hb expression can re-wire locomotor circuits, and it does so in three ways. First, it increases and re-distributes neuromuscular synapses. Second, it generates “hybrid” motor neurons that have axons, but not dendrites characteristic of early-born motor neurons. These two cell biological changes should lead to a change in the circuit level function of the U motor neuron temporal cohort. Third, prolonged expression of Hb reduces the number of neurons in the NB7-1 lineage with later-born molecular identities, which is likely to impact the uncharacterized circuits to which these neurons contribute. We see evidence that locomotor circuits are functionally re-wired because prolonged expression of Hb in specifically affects larval crawling behavior.

Thus, our data provides clear evidence that lineage-specific manipulation of single transcription factor is a potent regulator of circuit wiring. This observation provides insight into the types of molecular changes that could lead to the generation of novel circuits, either over evolutionary time scales or for biomedical intervention.

## Materials and Methods

### Fly genetics

Standard methods were used for propagating fly stocks. For all experiments, embryos and larvae were raised at 25°C, unless otherwise noted. The following lines were used: *CQ2-GAL4* (Bloomington stock center [BL] 7468), *OK6-GAL4* (BL 64199), *hsFLP; UAS(FRT.stop)myr∷smGdP-HA, UAS(FRT.stop)myr∷smGdP-V5-THS UAS(FRT.stop)myr∷smGdP-FLAG* (BL 64085), *UAS-myr-GFP* (BL 32198), *UAS-nls-GFP* (BL 32198), *UAS-Hb; UAS-HB*/*TM2* (BL 8504), *w1118* (BL 36005), *MHC-CD8-GCaMP6f-Sh* (BL 67739) *ac:VP16, gsb:v8v* (aka *NB7-1-GAL4*, gift of M. Kohwi).

### Tissue preparation

Three tissue preparations were used: Late stage whole mount embryos, in which antibody can still penetrate cuticle; isolated first instar (L1) CNSs, in which the CNS is removed from other larval tissue so that antibody reach the CNS; and third instar (L3) fillet preparations, in which the neuromuscular tissue and cuticle are dissected away from other tissue and pinned open like a book, allowing for superb immuno-labeling and visualization of larval neuromuscular synapses. For all preparations, standard methods were used for fixation in fresh 3.7% formaldehyde (Sigma-Aldrich, St. Louis, MO) [49-51]. For calcium imaging, L3 larvae expressing *MHC-CD8-GCamp6f-Sh* construct were dissected in HL3 solution containing 1.5mM Ca2+ and 25 mM Mg2+, brains removed, body walls rinsed fresh saline, and samples imaged.

### Immunotstaining

Tissue was blocked for an hour at room temperature or overnight at 4°C in phosphate buffered saline with 2% Normal Donkey Serum (Jackson ImmunoResearch), followed by 2 hours at room temperature in primary antibodies, and 1 hour at room temperature in secondary antibodies. Primary antibodies include: rabbit anti-Eve (1:1000, Heckscher lab, see below), chicken anti-GFP (1:1000, Aves #GFP-1020), chicken anti-V5 (1:300, Bethyl #A190-118A), mouse anti-HA (1:100, BioLegend #901501), rat anti-FLAG (1:300 Novus #NBP1-06712), rat anti-Worniu (1:250 Abcam #ab196362), goat anti-HRP-Cy3 (1:300, Jackson ImmunoResearch 123-165-021) rat anti-Runt (1:300, John Rientz, UChicago), guinea pig anti-Hunchback (1:1000, John Rientz, UChicago), guinea pig anti-Kruppel (1:1000, John Rientz, UChicago), rat anti-Zfh2 (1:800 Chris Doe, UOregon), rabbit anti-Castor (1:1000 Chris Doe, UOregon), guinea pig anti-Dbx (1:500 Heather Broiher, Case Western) guinea pig anti-HB9 (1:1000 Heather Broiher, Case Western). The following monoclonal antibodies were obtained from the Developmental Studies Hybridoma Bank, created by the NICHD of the NIH and maintained at The University of Iowa, Department of Biology, Iowa City, IA: mouse anti-myosin (1:100, 3E8-3D3), mouse anti-Futsch (1:50 22C10) mouse anti-Brp (1:50, NC82), mouse anti-DLG (1:500, 4F3), mouse anti-GluRIIA (1:25, 8B4D2), mouse anti-En (1:5, 4D9). Secondary antibodies were from Jackson ImmunoResearch and were reconstituted according to manufacturer’s instructions and used at 1:400. 647-Phallion (1:600 Jackson ImmunoResearch 123-025-021). Embryos were staged for imaging based on morphological criteria. Whole mount embryos and larval fillets were mounted in 90% Glycerol with 4% n-propyl gallate. Larvae brain preparations were mounted in DPX (Sigma-Aldrich, St. Louis, MO).

### Antibody Generation

An anti Even-skipped antibody was generated by GenScript (Piscataway, NJ). Rabbits were inoculated three times with a bacterial fusion protein containing an N-terminal 6xHis tag fused with the first 129 amino acids of Eve (Met…Arg Gln Arg). The resulting antiserum was immunopurified using the bacterial fusion protein.

### Single neuron labeling

Single U motor neurons were labeled by crossing a Multi-Color Flip Out fly line [52], harboring a heat-shock inducible FLP recombinase construct, to other lines of interest. For late stage embryo and L1 larval labeling, heat shock (37°C for 10 min) was delivered to embryos aged 6-24 hours on apple juice caps. For L3 labeling, late stage embryos and 1^st^ instar larvae were heat shocked and incubated at 25°C. All CNS tissue was co-stained with Eve antibody to confirm the identity of single cell clones. In Control, U MNs were identified by their characteristic position in the CNS with U1 positioned most medial and U5 lateral.

### Image acquisition

For fixed tissue images, data were acquired on a Zeiss 800 confocal microscope with 40X oil (NA 1.3) or 63 X oil (NA 1.4) objectives, or a Nikon C2+ confocal microscope with 40X (NA 1.25) or 60X (NA 1.49) objectives. For calcium imaging, data were acquired using a Zeiss 800 confocal microscope with a 40X dipping objective (NA 1.0) using 488-nm laser power with the pinhole entirely open. Images were acquired on a Zeiss 800 confocal microscope. Images were cropped in ImageJ (NIH) and assembled in Illustrator (Adobe).

### Image analysis

#### Cell body counting

We used *NB7-1-GAL4* driving *UAS-myristoylated-GFP* to count the number of cells within the NB7-1 lineage during late stage (16-17) embryos. Embryos were co-stained with Worniu and Eve antibodies to confirm that only the NB7-1 lineage clone was labeled. To ensure the entire lineage was labeled by GFP, we scored only segments in which both an Eve(+) U1 neuron (earliest born neuron in the lineage) and NB7-1 was labeled. Since the NB7-1 lineage clone is a dense cluster of cells in x-y-z, we used a custom Fiji plug-in, that employs the *Multi point* tool to count cells.

#### Type 1b branch counting

We stained against HRP to detect the neuronal membrane and Discs large (Dlg) in L3 fillet preparations. Dlg allowed us to distinguish between type 1b and type 1s boutons. We counted contiguous stretches of Dlg staining that overlapped with HRP as a single branch.

#### Calcium imaging

X-y-z-t stacks (Figure S3B-C) were converted into x-y time series images using the Maximum Intensity projection function (Fiji). X-y time series images were then registered using the Register Virtual Stack Slices plug-in (Fiji) to reduce movement artifacts. Time series images were projected into two different x-y single images using either the Maximum Intensity projection function (Fiji) or Average Intensity projection function (Fiji), and then the average intensity was subtracted from the maximum to get a change in fluorescence image (Figure S3D).

### Statistics

Descriptive statistics: average and standard deviation are reported, except for behavior, where the average of average speed and standard error of the mean are reported. Every data point is plotted in figures. Test statistics: All data was assumed to follow a Gaussian distribution. If standard deviations were unmatched Welch’s correction was applied. For numerical data in two populations, we used unparied, two-tailed t tests. For numerical data in more than two populations, we used ordinary one-way ANOVAs with Dunnett or Games-Howell correction for multiple comparison. Analysis done using GraphPad Prism.

### Larval Behavior

The behavioral set-up was described previously [24]. Briefly, 10-50 larva were allowed to crawl on a 2% agarose arena for at least 10 min prior to recording. The arenas were placed on a 7.5 diameter circle of plexiglass illuminated with 850 nm LEDs (1 m of SMD3528-600 LED light strip; 120 LEDs per meter at 9.6W/meter; HK-F3528IR60-X from ledlightsworld.com), in a modified frustrated total internal reflection set-up ([24]). Behavior was recorded with Point Grey FlyCap2 software and attached to a GS3-U3-41C6NIR camera (Point Grey) mounted with a 16mm Standard Schneider Compact VIS-NIR Lens and 825 nM high performance long-pass filter (Edmund Industrial Optics, 86-070). Images were acquired at 10 frames per second.

## Acknowledgements

N. Grace Schulz, Machiah Gill, Catarina Machado De Oliveria Catela, Xiaoxi Zhang, Marie Greaney, Edwin Ferguson, Sally Horne-Badovinac, 2018 Drosophila Neurobiology: Genes, Circuits & Behavior course, T32 GM007183 to JM and NIH R01-NS105748, MGCB start-up funds to ESH.

---

**Figure S1.**
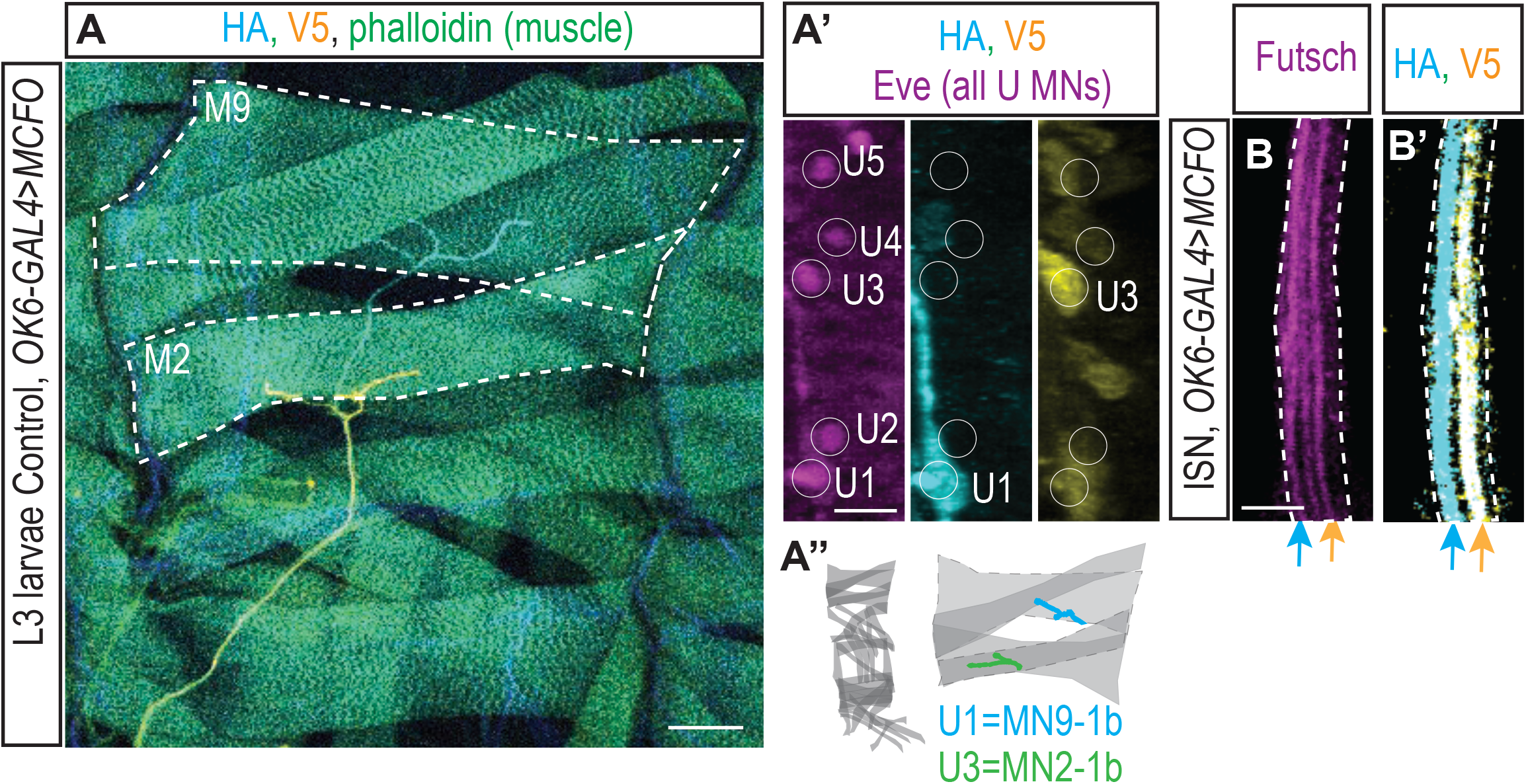
Single neuron labeling in Control larvae. (A-A’’) Images show example of single motor neurons labeling in Control L3 larva. Using the pan-motor neuron driver, OK6-GAL4 we drove MCFO transgenes (*hsFlp; OK6 GAL4/+; UAS(FRT.stop)myr∷smGdP-HA, UAS(FRT.stop)myr∷smGdP-V5-THS-UAS(FRT.stop)myr∷smGdP-FLAG/+).* The dorsal muscle field shows MN9-1b labeled by HA and MN2-1b labeled by V5 (A). In the CNS, HA labels U1 and V4 labels U3. (A’’) Illustration of representing labeling in A’, with all muscles shown at left and labeled neuromuscular synapses overlaid at right. (B-B’) Images of the intersegmental nerve (ISN) over Muscle 3 in Control. One Futsch-labeled bundle (B) runs through each individually-labeled MN (B’). Borders of the ISN are shown (dashed lines). Blue and yellow arrows point to same neuron in B and B’. Scale bars represent 50 microns (A) and 5 microns (A’, B, B’).

**Figure S2.**
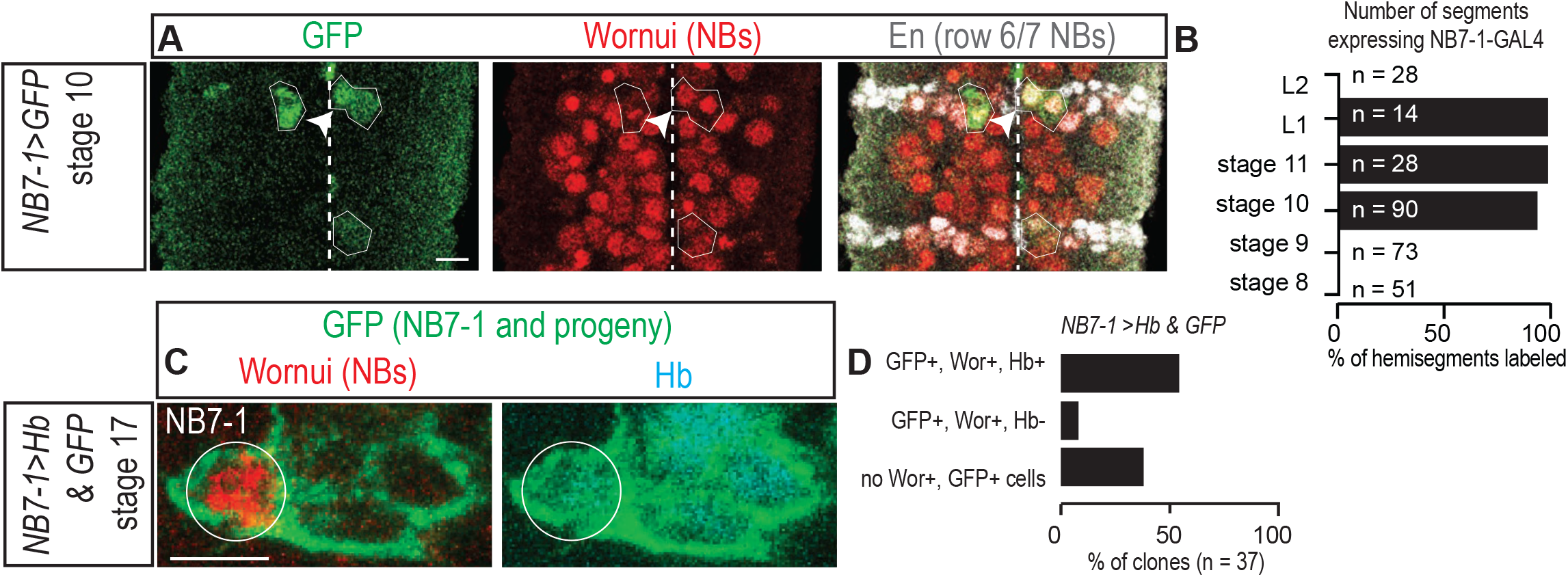
Characterization of *NB7-1-GAL4*. (A) Image of *NB7-1-GAL4* expression in Control embryos. *NB7-1-GAL4* is driving GFP expression (*gooseberry-DBD, achaete-VP16/+; UAS-nls-GFP/+*), all row 6 and row 7 neuroblasts are labeled with En (white), and all neuroblasts with Wor (red). Arrowhead points to NB6-1, which is occasionally labeled by *NB7-1-GAL4*. One complete segment is shown with anterior up, and midline dashed. (B) Quantification of percent of hemisegments expressing GFP under the control of *NB7-1-GAL4* at different stages. (C) Image of a late stage embryo showing NB7-1 and its latest born progeny in NB7-1>Hb. NB7-1 expresses a neuroblast marker (Wor), GFP, and Hb. (D) Quantification of Hb expression in NB7-1 in late stage embryos. In a majority of segments, there are GFP(+) Wor(+) Hb(+) cells, showing Hb is expressed throughout neurogenesis in NB7-1. In fewer segments, there are GFP(+) Wor(+) Hb(-) cells, or no cells expressing GFP and Wor. Images are shown with anterior is up. Scale bars represents 5 microns.

**Figure S3.**
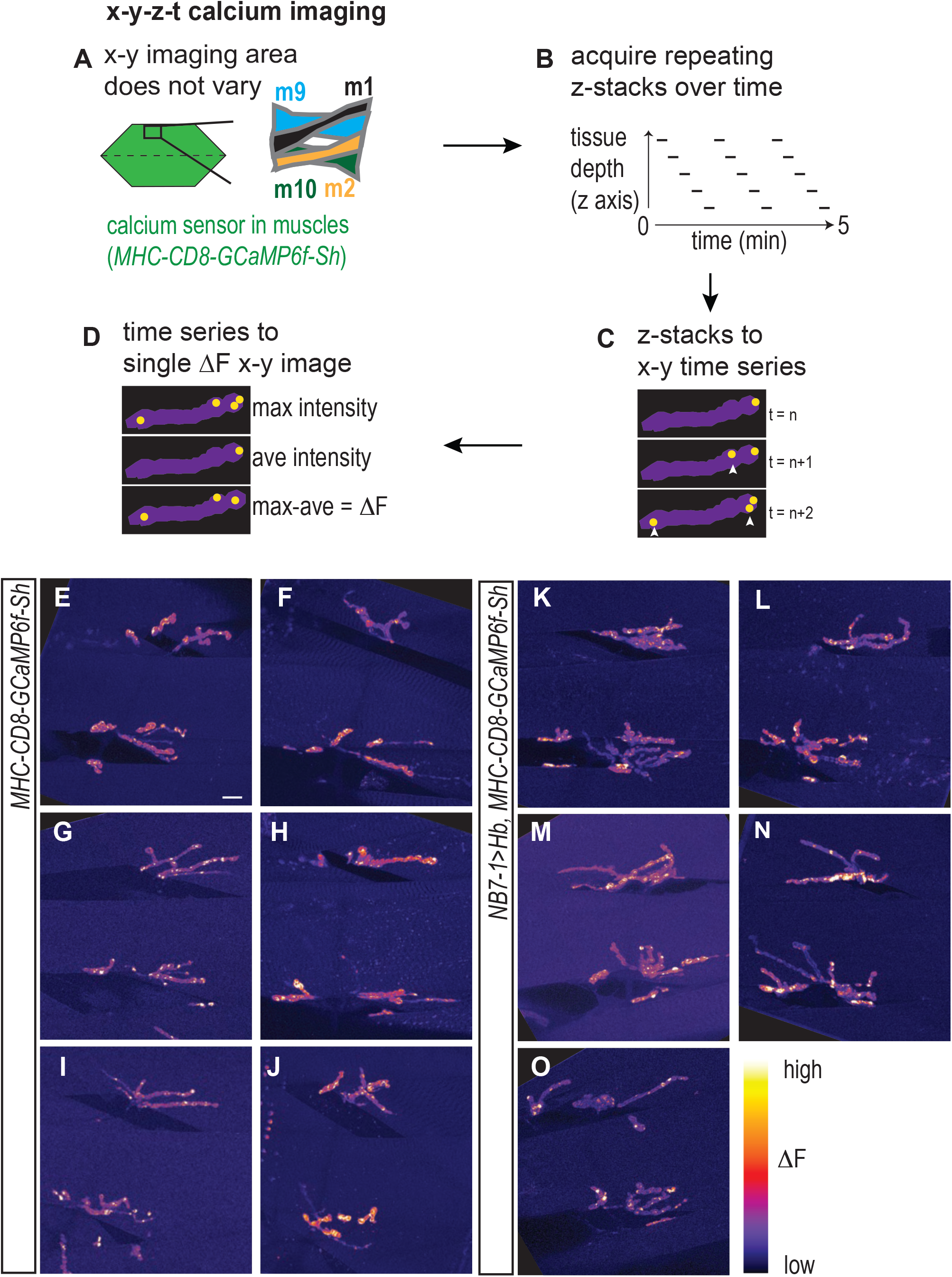
Calcium imaging protocol, analysis, and additional examples. (A-D) Illustration of calcium imaging protocol and analysis are shown. See Methods for details. (E-O) Images of calcium signals on Muscles 1, 2, 9, 10 for Control (*NB7-1-GAL4/+; MHC-CD8-GCaMP6f-Sh/+)* and NB7-1>Hb (*NB7-1 GAL4/UAS Hb; MHC-CD8-GCaMP6f-Sh/UAS Hb)*. Scale bar represents 5 microns.

**Figure S4.**
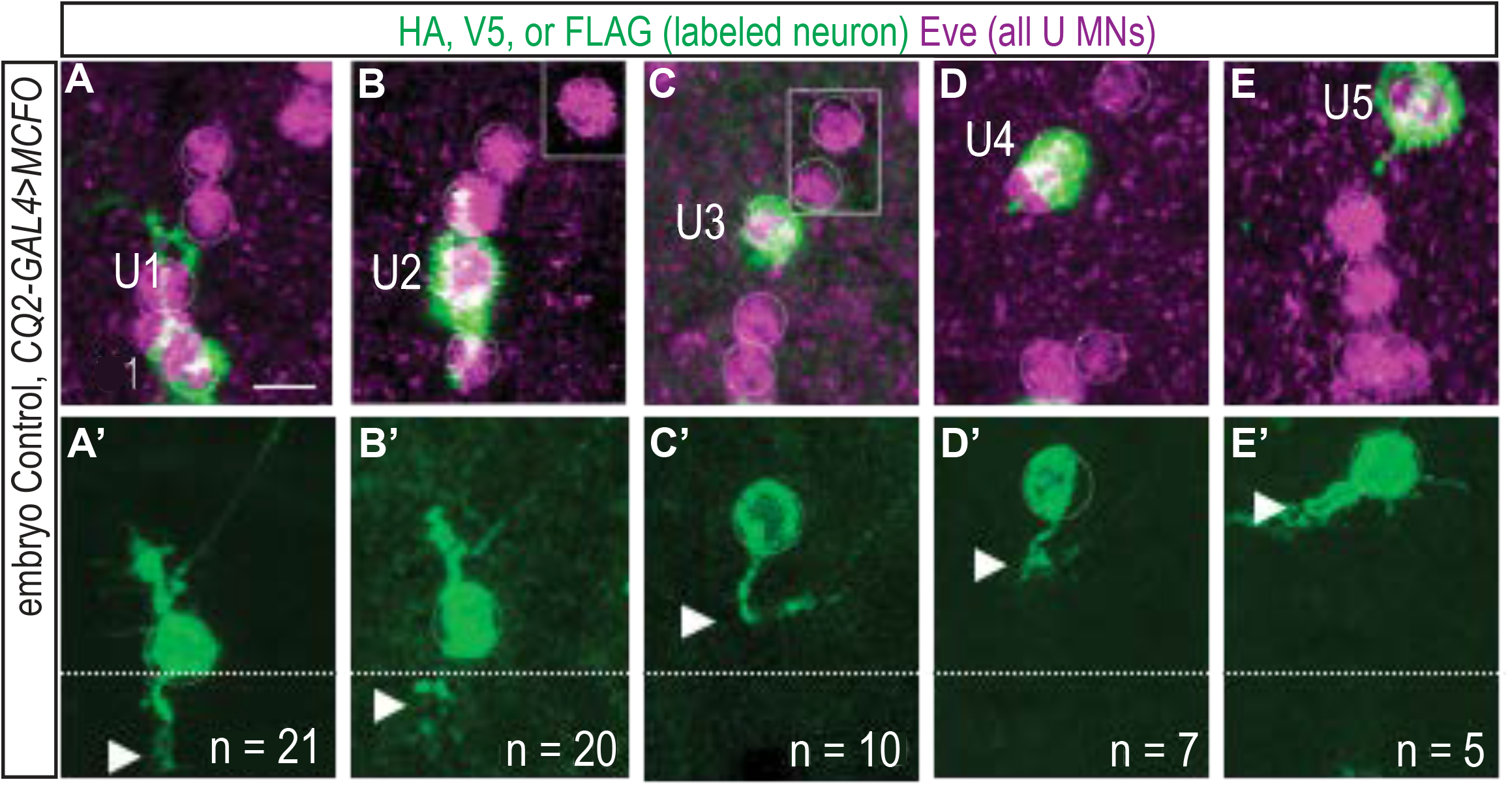
Single neuron labeling in Control larvae. (A-D, A’-D’) Image of the late-stage embryonic CNS 17 in Controls. Individual U motor neurons were labeled (*hsFlp; CQ2-GAL4/+; UAS(FRT.stop)myr∷smGdP-HA, UAS(FRT.stop)myr∷smGdP-V5-THS-UAS(FRT.stop)myr∷smGdP-FLAG/+).* U1 and U2 send dendrites across the midline (dashed), and U3-U5 do not send dendrites across the midline.. Arrowheads point to the distal-most dendrite tip. Boxes are insets from other focal planes. Number of single neurons of each type that were labeled is shown. These images are of the same neurons as those in Figure 1D-H’. Scale bar represents 5 microns.

